# Stochasticity in bacterial division control: Preliminary consequences for protein concentration

**DOI:** 10.1101/826867

**Authors:** Cesar Augusto Nieto Acuna, Cesar Augusto Vargas Garcia, Juan Manuel Pedraza

## Abstract

The stochastic nature of protein concentration inside cells can have important consequences in their physiology and population fitness. Classical models of gene expression consider these processes as first-order reactions with little dependence with the cell size. However, the concentrations of the relevant molecules depend directly on the cellular volume. Here we model the cell size dynamics as exponential growth followed by division with occurrence rate proportional to the size. This framework, together with known models of chromosome replication and both protein and mRNA synthesis, lets us predict relationships between cell size and both protein number and concentration. As a main result, we find that protein production strategies (constant rate or rate proportional to either chromosome number, cell size or chromosome number times cell size) can be experimentally distinguished from the correlation between protein concentration and cell size.

## INTRODUCTION

Stochastic modeling for molecular biology has been successful explaining some important phenomena such as variability in fluctuations of protein concentration at single-cell level (Elowitz, et al. 2002, Kaern, et al. 2005, Blake, et al. 2003), ion channel gating (Smith 2002, Monod, Wyman and Changeux 1965), bacterial chemiotaxis (Falcke 2004), intracellullar transport (Bressloff and Newby 2003) and self-organization (Misteli 2001). Some of these predictions have been challenged by increasingly detailed experiments (Mogilner, Wollman and Marshall 2006, Duncombe, Tentori and & Herr 2015).

Stochastic effects are important since in living cells many components are present at low copy numbers (mRNA, DNA loci, some transcription factors, etc) (McAdams 1997) and thus, their fluctuations could be relatively high. Once cellular components interact with one another in complex regulatory networks, some of them non-linearly, this variability is amplified and converted into a distribution of phenotypes, explaining cell-cell variations in clonal populations (Blake, Balázsi, et al. 2006).

Many consequences of this heterogeneity have been studied (Raj and van Oudenaarden 2008). Benefits include facilitating evolutionary transitions (Eldar and Elowitz 2010), increasing the population fitness in a fluctuating environments (Kussell and Leibler 2005, Garcia-Bernardo and Dunlop 2016), improving the bacteria stress response (Süel, et al. 2007) and surviving through sudden antibiotic encounters (Balaban, et al. 2004).

Cell division is suspected to explain part of this fluctuations in gene expression (Amir 2014, Modi, et al. 2017). In rod shaped organisms, physiological implications can potentially affect DNA concentration, surface transport and biosynthesis rates, as well as proteome composition (Willis and Huang 2017). Current models for cell division try to predict some characteristics of bacterial cell size such as size distributions (Koch 2001), effects of fluctuations in division control in terms of population fitness (Hashimoto, et al. 2016) and auto-correlation and spectral analysis of division strategies through several generations (Tanouchi, et al. 2015).

In this paper, we present a stochastic model for describing the cell division as a continuous rate Markov chain (CTMC) (Davis 1984, Gardiner 1985) where the number of divisions defines the states of the CTMC and the transition rate between these states is proportional to the current size of the bacteria. We discuss its physiological implications for the statistics of protein concentration.

## MATERIALS AND METHODS

### The classical model for stochastic gene expression

Stochastic models for bacterial cell physiology usually describe the dynamics of quantitative variables like cell size, protein counting and protein concentration as random variables modeled by continuous time Markov chains (De Jong 2002). The protein synthesis-degradation, specifically, is simplified as a Birth-Death process (Paulsson 2005). By this, protein synthesis is modeled with the protein synthesis rate (*k*_*c*_) as a first order chemical reaction. This rate is a function of the concentration of control molecules called Transcription Factors (*T*). Degradation rate (*γ*) is proportional to the current concentration (*c*) of the given protein. Protein concentration (*c*) is represented by

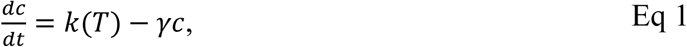

where we define *k*(*T*) = *γc*_0_(*T*) with *c*_0_ as the steady protein concentration.

If mRNA dynamics are included, Eq 1 turns into a system of differential equations where protein synthesis is proportional to the number of mRNA and these are transcribed at a rate proportional to the TF’s concentration. However, taking advantage of the shorter life-time of mRNA compared to proteins a “translational bursting” approximation is taken (Shahrezaei and Swain 2008). This assumes that effects in the noise due to mRNA transcription can be approximated if a burst of proteins is translated through the time-life of the mRNA molecule. Later, some additional details will be given.

Constitutive proteins are produced by the cell continuously and its concentration, on average, is kept constant robustly through time. Robustness is usually obtained considering the production rate as a zero-order chemical reaction, ie., *k*(*T*) = *k*_*p*_ (Thus, *c*_0_ = *k*_*p*_/*γ*).

A heuristic way to obtain Eq 1 is to assume that proteins are produced at a volume-dependent rate. If *p* is the number of proteins inside a cell of size *s* then average protein count dynamics can be written as

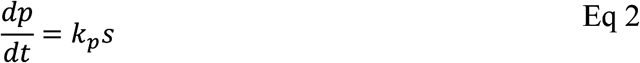

Defining the concentration 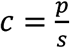 and assuming that cell size grows exponentially at rate 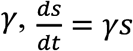, Eq 1 yields:

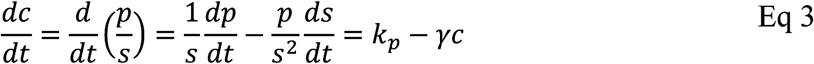

Traditionally, fluctuations are added to this by a white-noise term (*η*) with intensity *q*

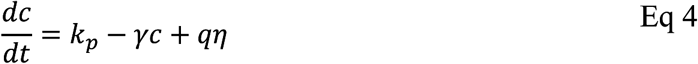

Where white-noise is a random variable with mean zero, unitary variance and uncorrelated with itself at other times, i.e., ⟨*η*(*t*), *η*(*t*′)⟩ = *δ*(*t* − *t*′) with *δ*(*x*) the Dirac delta distribution. The intensity of the noise, *q*, can be obtained analytically for the intrinsic noise of creation and destruction with constant rates but is usually obtained from experiments as an effective parameter for other sources of noise.

Advantages of this model include its linearity, yielding analytical formulas for the distribution of protein concentration (Poisson and super-Poisson distributions, for instance), with simple analytical expressions describing the statistical moments. However, this approach excludes some fundamental biological details of bacteria physiology like division effects (Huh and Paulsson 2011), chromosome segregation (Reyes-Lamothe, Nicolas and Sherratt 2012) and size control strategies (Taheri-Araghi, et al. 2015). Transition rates and stochastic simulation algorithms (SSA).

Many detailed stochastic models for gene expression see it as a counting process. This kind of processes describe the occurrence of events. These events occur at a given rate *k*. This means that the probability of occurrence of this process during an infinitesimal time interval (*t, t* + *dt*) is:

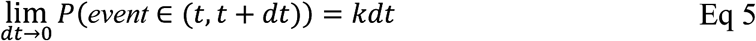

The rate *k* is not generally constant and is also called propensity function or hazard function. Through integration of Eq 5, let us obtain the probability of the event not happening during the time interval (0, *t*) and happen during (*t, t* + *dt*):

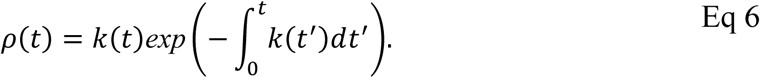

In the particular case of *k*(*t*) = *k*_*p*_, a constant, we see that *ρ*(*t*) is an exponential distribution. This kind of processes are called “memoryless”: the probability of occurrence in an infinitesimal time interval (Eq 5) is not dependent on the time.

### Division strategies and chromosome segregation

How bacteria decide to divide while maintaining their size in a narrow distribution (the standard deviation over the mean (CV) is approx. 14%) (Taheri-Araghi, et al. 2015) and how the division is synchronized with chromosome replication are long standing problems (Cooper and Helmstetter 1968, Schaechter, Maaløe and Kjeldgaard 1958) that are currently not fully understood.

Current models describing division propose two mechanisms acting independently: the division and the chromosome replication (Si, et al. 2019). Some efforts understanding division mechanisms have been done both theoretically (Amir 2014, Ghusinga, Vargas-Garcia and Singh 2016) and experimentally (Taheri-Araghi, et al. 2015, Campos, et al. 2014). Division has been studied by quantifying the relationship between size at division and the size immediately after the last division (or, simply, size at birth). No correlation was found between the added size and the size at birth, which is unexpected given some classical division strategies. If cells divide once a specific size is reached negative correlation is expected and if cells wait a specific time before division, positive correlation is expected.

This decorrelation between size at birth and added size defines the called “Adder” mechanism found in some different bacteria like *E. coli* (Taheri-Araghi, et al. 2015), *B. subtilis* (Campos, et al. 2014), *C. crescentus* (Iyer-Biswas, et al. 2014) and in some archea (Eun, et al. 2018). In *E. coli* underlying mechanism seems to be related to the formation of the FtsZ ring which triggers division (Si, et al. 2019).

Chromosome replication has also been studied experimental and theoretically (Wallden, et al. 2016, Levin and Taheri-Araghi 2019). The current paradigm, also called “initiation adder” (Wallden, et al. 2016) states that chromosome replication starts once cell has reached a fixed “size per origin” allowing multiple replication “forks” if bacteria size is large enough to have more than one “size per origin”. The time to complete the chromosome replication is nearly constant (C in Figure 1); if growth media is rich enough to let bacteria duplicate with generation times faster than this time (C), the largest bacteria can have more than one replication happening simultaneously.

**Figure 1.**
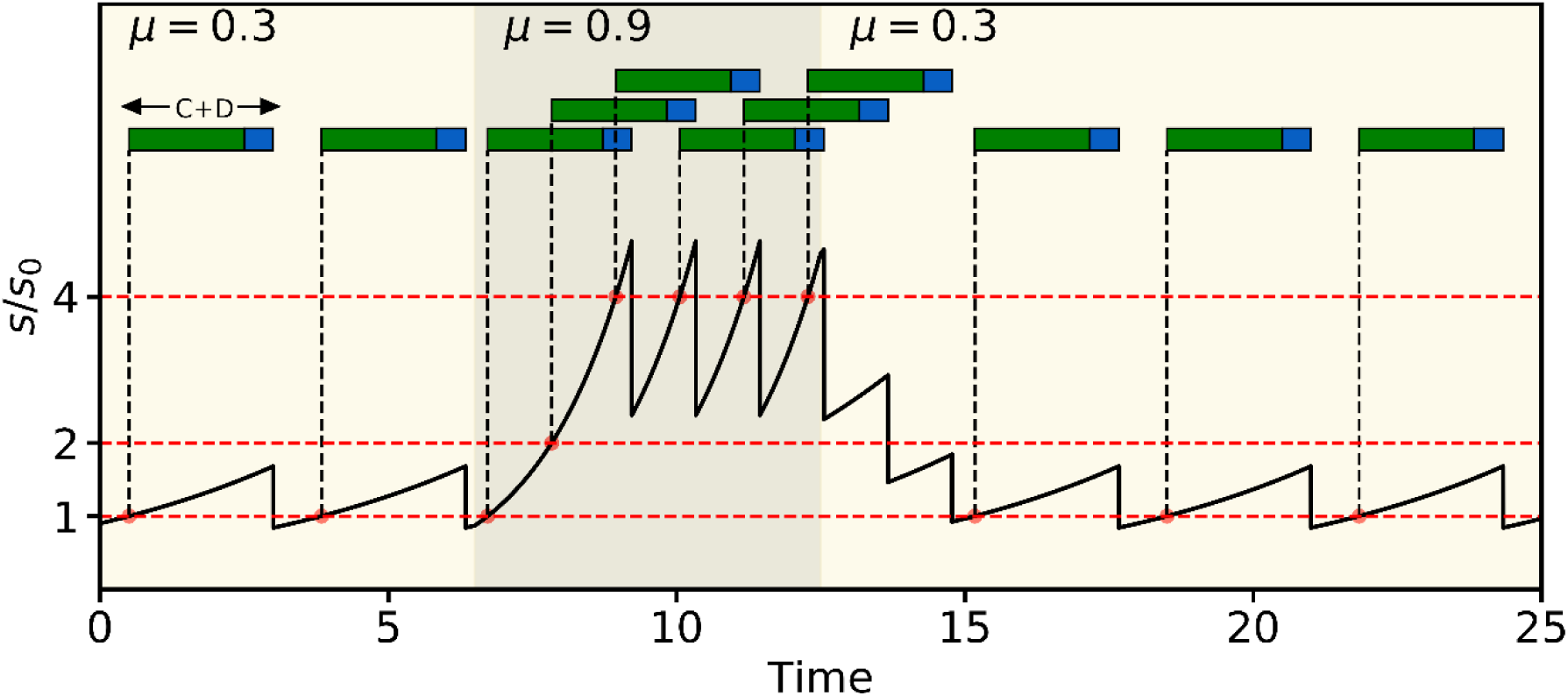
Simulated volume expansion and division for an idealized cell lineage going through an upshift and a downshift in growth rate. Replication is initiated at a fixed volume per chromosome (red circle), and the cells divide a fixed period of time later, including the required time for replication (C-period) and chromosome segregation and septum formation (D-period). A dashed green arrow indicates the relation between initiation of replication and its corresponding division. Adapted from (Wallden, et al. 2016).

The mechanism explaining how bacteria know if the current size is big enough to start the duplication is presumed to be related to multiple systems regulating oriC and DnaA genes (Skarstad and Katayama 2013). As can be seen in Figure 1, the inclusion of a D period after the chromosome duplication period (C) coincides with the idea of the division being triggered not by the chromosome replication but by an additional mechanism as the theory of division adder predicts.

Figure 1 illustrates an important point: the copy number of genes changes at some point along the cell cycle. The timing and total number depends on the position of a gene along the chromosome, but in some cases the copy number can be as high as 8 when a third replication has started. This gene copy number variation can impact the noise in gene expression in ways that classical models are unable to capture.

## RESULTS

### Division using a continuous rate model

Consider a cell with size *s*_*b*_ after last division, which is growing exponentially in time at rate *μ*. Now we assume that divisions are occurring stochastically at rate proportional to the size. This is:

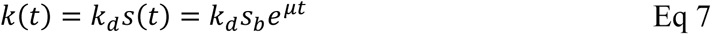

Integration of Eq 7 over time following Eq 6 gives us the probability density distribution of cell division (*t*_*d*_)

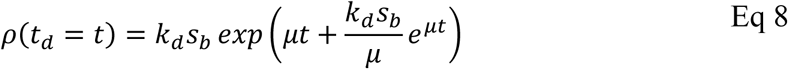

Which, using the transformation 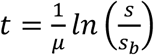, after a change of variables, gives us the probability density of the division size *s*_*d*_ being *s*:

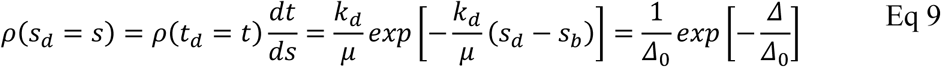

which corresponds to an exponential distribution of *Δ* = *s*_*d*_ − *s*_*b*_ with means 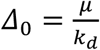, which in turn defines the expected values: ⟨*s*_*d*_⟩ = 2*Δ*_0_; ⟨*s*_*b*_⟩ = *Δ*_0_.

The size at division (*s*_*d*_) is a function of the added size (*Δ*) and this is independent of the size at birth (*s*_*b*_), so this model is a simple *adder* as was experimentally found (Campos, et al. 2014). However, the exponential distribution (with *CV* = 1) obtained is not similar to the experimental findings (*CV* ≈ 0.1). We already discussed how to overcome this situation considering the occurrence of not only one event but more than one with similar rate proportional to the size (Nieto-Acuna, et al. 2019). Including details of this idea would make the model more complex and could overshadow the results of the following calculations.

Knowing the transition rate and its dependence on the time given by Eq 7, we can use the model to predict the division dynamics and describe the protein concentration in growing cells. Unlike the classical model, using our framework, one can predict some interesting properties like the correlation between protein concentration and cell size.

### Protein expression models

As we explained above, gene expression has been studied as a first order chemical reaction and Eq 1 can be obtained considering a protein synthesis with rate proportional to the current cell size (Eq 2). However, there is not, a priori, an argument for this to be the general case. Thus, we will explore some additional possible cases to examine their possible signatures. An experimental verification of these would be an important argument for choosing one of the possible hypotheses.

We will consider the division strategy described by the division rate given by Eq 7 and explore how protein synthesis may change following different strategies where the production rate is given by equation 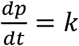 with *k* being the synthesis rate given by one of the following strategies:

- **A constant protein production rate (*k*_*p*_):** The protein production rate is not dependent on the size but simply constant. Biologically, this can be seen as a production dependent on molecules whose amount is held fixed and therefore their concentration is inverse to the cell size.
- **Size dependent protein production rate (*k*_*p*_*s*):** Protein production rate proportional to the size as Eq 2 and classical model assume. Biologically this model can be seen as protein production due to molecules whose concentrations are held constant independently of the cell size.
- **A protein production rate proportional to the number of chromosomes (*k*_*p*_*n*):** Chromosome is duplicated at rate proportional to the size, similarly to Eq 7, starting once the cell has reached the minimum size per origin *s*_0_ = 0.8*Δ*_0_ following the strategy depicted in Figure 1. Then, we assume the rate of protein synthesis is proportional to the number of chromosomes assuming values in {*k*_*p*_, 2*k*_*p*_, 4*k*_*p*_, …} depending on the value of *n*. Biologically this would be a condition where the DNA is limiting.

**A protein production rate proportional to the number of chromosomes times the size (*k*_*p*_*n*s):** Similar to the (*k*_*p*_*n*) model but in this case, the protein production rate is not only proportional to the number of chromosomes but also to the size as classical model assumes. Biologically this would be a condition where both the DNA and molecules like polymerases are limiting.

### Statistics of protein number

As we showed, the dynamics of concentration, Eq 3, are naturally derived from the model of protein synthesis, Eq 2, considering a protein synthesis rate proportional to the size and dilution due to growth. We now compare the four possible protein production including multiple stochastic processes:

- **mRNA transcription:** This process is an intermediate process before protein synthesis. We modeled it as a counting stochastic process with production rate given in turn by each of the four models of synthesis under study. The magnitude of these rates was fixed such that the mean mRNA concentration would be 10 mRNA molecules per cell. Every time the cell divides, if the number of mRNA is even, the cell keeps the half of this number. If the number is odd, the cell keeps with equal probability either the higher or lower integer closest to half of the number of mRNA before division. No active degradation was considered, although including a degradation faster than protein dilution could add a reduction on the noise due to time averaging.
- **Protein synthesis:** We consider a production rate proportional to the number of mRNA inside the cell as the classical model does. We choose the rates such that the main concentration of proteins would be 1000 and, during the division, protein numbers split in the same way as mRNA.
- **Division**: A Markov process is considered with occurrence rate proportional to the current size (Eq 7). The distribution of times of division given the size at birth is given by Eq 8. During the division, we considered perfectly symmetric division (cell splits on one half perfectly). Including well studied (Huh and Paulsson, 2011) models of noise in cell partitioning would increase the noise.
- **Chromosome duplication**: Depending on the production strategy taken, chromosome duplication is present or not. In the cases where it is present, we consider a chromosome synthesis following the Helmstetter & Cooper model (H&C). Here the bacterial cell could have a number of chromosomes *n* ∈ {1,2,4 … 2*m*}. During division cell keeps the half of this number. Synthesis rate is proportional to the size as given by:

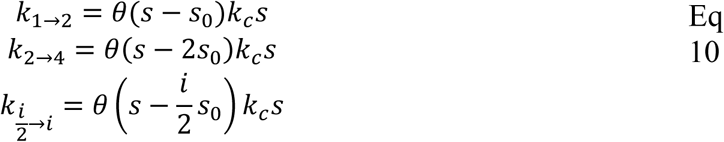
- With *θ*(*x*) the Heaviside step function. The main parameters of this process are the size per origin (*s*_0_) and the sinthesys rate (*k*_c_) with these values being constant independently of the growth rate (Wallden, et al. 2016). Here we take as main units of size and chemical kinetics the mean added size 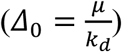 in Eq 9 and the division rate *k*_*d*_. Thus, we can fix *s*_0_ and *k*_c_ in order to have either a fast or a slow growth. In fast growth, the cell size is relatively large and *s*_0_ is small. Hence, we use *s*_0_ = 0.6*Δ*_0_ and the chromosome replication is then relatively slow with respect to the division rate. Because of this, we set *k*_c_ = 0.7*k*_*d*_. To explore the statistics of protein level, we use the fast growth parameters since this is one of the most studied regimes (Taheri-Araghi, et al. 2015).

### Stochastic Simulation Algorithm

To simulate a population of independent cells our procedure will be detailed as follows: Define the number of cells to simulate. We took 5000. Every time these cells divide, we keep only one descendant such as the number of cells is constant during the simulation.

1. Every cell has certain properties: Size, number of chromosomes, number of mRNA and number of proteins. Rates of synthesis, replication and growth are given by the models explained above.
2. Define a time step Δ*t* ≪ *k*_*p*_^−1^ ≪ *μ*^−1^. Calculate the probability of occurrence of all the stochastic events during this Δ*t* by using the occurrence rates *h*(*t*′) and integrating equation

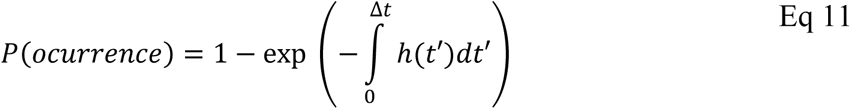
3. Generate a random number (*R*) uniformly distributed in (0,1). If this number is less than *P*(*ocurrence*), the event happens. This is done for all the chemical reactions considered in cell cycle: Division occurrence, mRNA transcription, DNA translation and chromosome duplication.
4. Deterministic variables evolve. For instance, the time *t* = *t* + Δ*t* and cell size *s* =*s* exp(*μ*Δ*t*).
5. Repeat steps 3,4 and 5 for all the cells until the maximum time of simulation (*t* = *t*_*max*_) is reached. We took this time to be 10 division times approximately such that all the moments of the distributions of sizes and protein numbers reach values approximately constant.
6. Statistics of the number of proteins

Histograms of the number of proteins inside each cell are shown in Figure 2. These distributions correspond to histograms obtained simulating 5000 cell using a Montecarlo simulation. All of them have mRNA transcription, stochastic division and protein synthesis as explained before. Cells with protein synthesis dependent on the number of chromosomes have a stochastic version of the H&C model.

**Figure 2.**
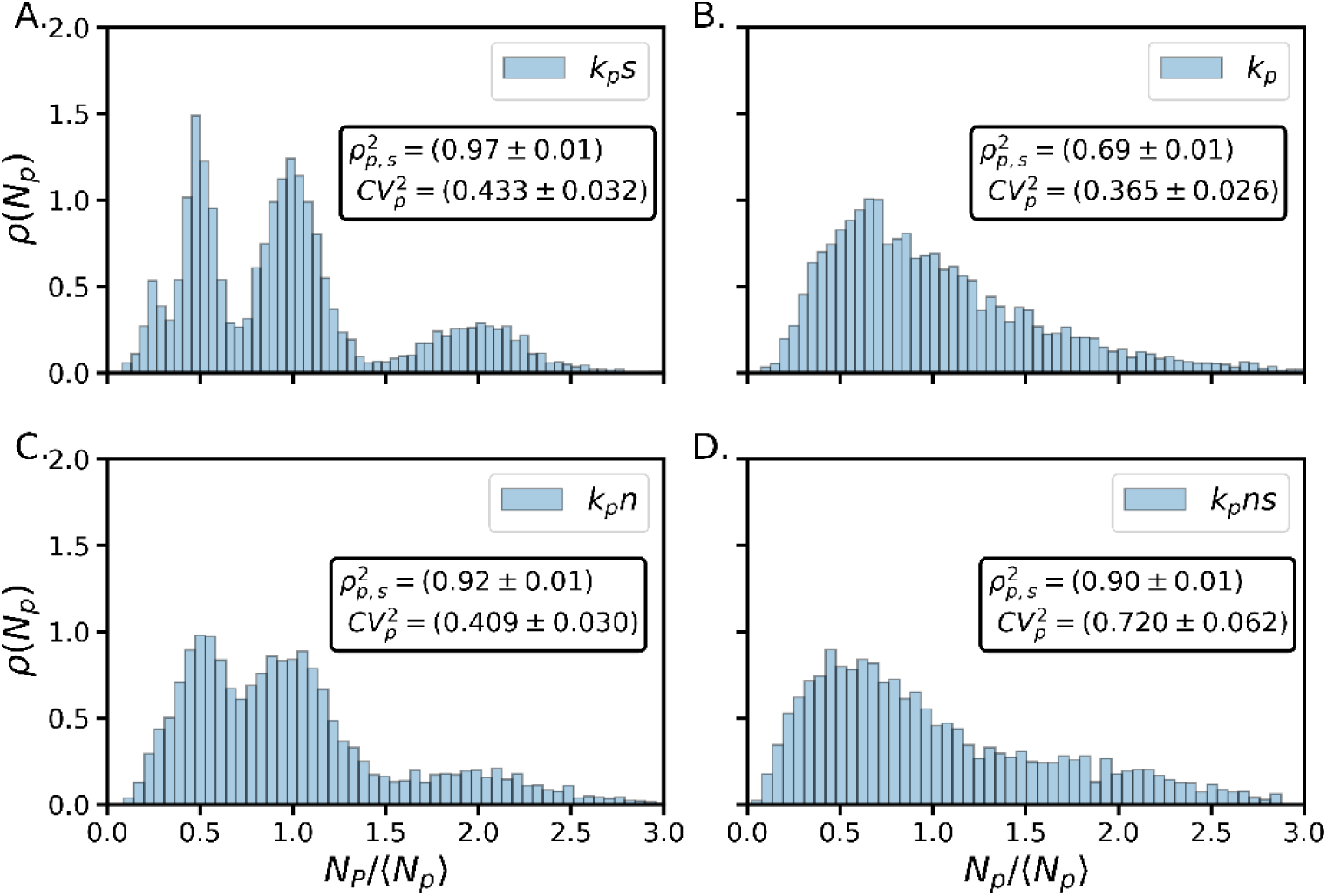
Stationary distribution of protein number obtained using different protein synthesis strategies. A. Size dependent protein production rate (*k*_*p*_*s*) B. Constant protein production rate (*k*_*p*_). C. Protein production rate proportional to the number of chromosomes (*k*_*p*_*n*): D. Protein production rate proportional to the number of chromosomes times the size (*k*_*p*_*ns*).

In figure 2, boxes show the Pearson correlation coefficient between the protein number and cell size showing a high correlation independently of the particular strategy. This high correlation leads to multimodality in some distributions, for instance, in the size dependent synthesis rate strategy. Each peak in the distribution correspond to bunches of proteins with cell sizes proportional to a power of 2 times the mean size. This is 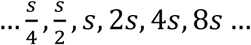 and due to this correlation, we see peaks centered in protein number proportional to 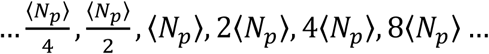

### Statistics of protein concentration

Protein concentration, in addition to the cell size, is perhaps the variable that can be measured most easily, usually by tagging with fluorescent proteins whose fluorescence intensity can be measured using microscopy imaging and assuming that the intensity of the fluorescence is proportional to this concentration. The concentration distribution for different strategies of protein synthesis is shown in Fig 3.

**Figure 3.**
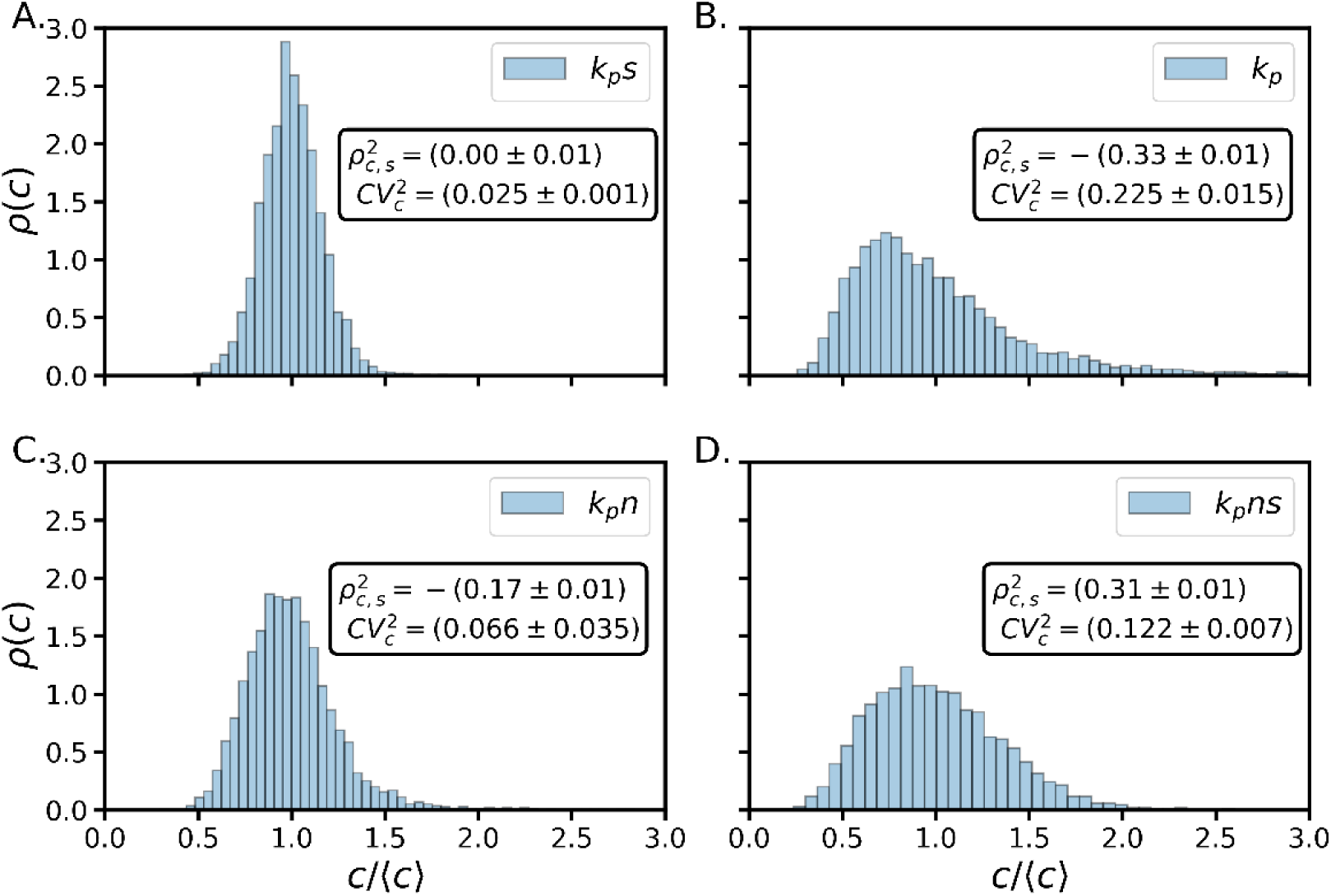
Stationary distribution of protein concentration obtained using different protein synthesis strategies. A. Size dependent protein production rate (*k*_*p*_*s*) B. constant protein production rate (*k*_*p*_). C. Protein production rate proportional to the number of chromosomes (*k*_*p*_*n*): D. Protein production rate proportional to the number of chromosomes times the size (*k*_*p*_*ns*).

Unlike the distribution of number of proteins, protein concentration is unimodal and with a correlation with cell size that depends on the production strategy. The correlation between size and protein concentration could be negative (*k*_*p*_and *k*_*p*_*n*), zero (*k*_*p*_*s*) or positive (*k*_*p*_*ns*). The different possibilities for this coefficient let us propose it as parameter to distinguish between strategies of protein production.

## DISCUSSION

In this work, we propose a model that describes de division of rod-shaped organisms based on recent studies (Vargas-García and Singh 2018) and observations (Si, et al. 2019). This approach considers the division triggered by a stochastic process with occurrence rate proportional to the size.

Dependence with the size of this splitting rate function can be related to the synthesis of a protein which number triggers the division. This idea has been explored in some studies (Ho and Amir 2015, Ghusinga, Vargas-Garcia and Singh 2016, Chandler-Brown, et al. 2017., Delarue, Weissman and Hallatschek 2017). If zero order chemical production is considered, protein number inside the cell follows the Eq 2, which is exactly our splitting rate. All free parameters of the division model can be estimated from observations. For instance, *k*_*d*_ can be estimated measuring the average cell size (proportional to 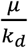) without need for additional or more complicated experiments.

## CONCLUSIONS

In this work, we propose a model that describes de division of rod-shaped organisms based on recent studies and observations describing cell division, protein synthesis and chromosome replication. This approach considers cell division to be triggered by a stochastic process with occurrence rate proportional to the size. We use Stochastic Simulation Algorithms to estimate distributions and correlations for different possible models of gene expression. We found that the stationary distribution of both concentration and number of proteins can be obtained together with the dependence of these variables on the size. Our results reveal that protein production mechanisms can be distinguished based on the correlation between protein concentration and cell size.

Applications of this framework are wide. Modeling gene expression dynamics and protein concentration inside the cell is an important problem in molecular biology (Paulsson 2005). Our theory can be used to explore in more detail the dependence of these molecular phenomena including the size of the cells. A particular advantage is that it does not identify the specific molecules so it can apply to multiple organisms, and the small number of parameters allow it to work also as a phenomenological description in cases where the biological details are still unknown.

## AKNOWLEDGEMENTS

We thank COLCIENCIAS convocatoria para doctorados 647 for financial support.

